# Status of the pillar coral *Dendrogyra cylindrus* in Los Roques National Park, Southern Caribbean

**DOI:** 10.1101/2020.09.15.297770

**Authors:** F. Cavada-Blanco, J. Cappelletto, E. Agudo-Adriani, S. Martinez, JP. Rodriguez, A. Croquer

## Abstract

Information on the status of the pillar coral *Dendrogyra cylindrus* across its global distribution range is needed to plan and implement effective conservation interventions at both the national and regional level. Knowledge on the species distribution and abundance on the southernmost edge of its range was limited to qualitative data gathered in the 1980s. In 2014, we started using local ecological knowledge and extensive surveys to assess the status of the pillar coral in Archipelago Los Roques National Park, Venezuela; also evaluating the species representativeness within the MPA according to the level of protection established by the park’s zoning. Between 2014 and 2016, we recorded over 1,000 colonies-the highest abundance reported to date for the species- within 14 different habitat types. Disease, bleaching and partial mortality prevalence were below 4%. Size frequency distribution was uni-modal for the MPA and dominated by medium size colonies (40cm -70cm height) suggesting potential for intrinsic population growth. However, the structure of size classes varied among reefs (Pseudo-F=2.70, p=0.03), indicating asynchronous dynamics mostly driven by reef-scale processes. Overall, our results indicate that Los Roques could be a stronghold for the species. But, to maintain the conservation value for coral reefs and the pillar coral, the MPA’s zoning designation needs to be urgently revised and the extension of its high-protection zones expanded to increase habitat redundancy as well as the singular habitats composed by thickets of *Acropora cervicornis* and mounds of *Madracis* sp. This work confirms the species as extant in one of the four localities within its national range in Venezuela. However, further research on genetic diversity and connectivity among reefs within the MPA is needed to estimate effective population size and assess viability.

## INTRODUCTION

According to the last global Red List assessment for Scleractinian corals, one third of assessed Caribbean species are threatened with extinction (Carpenter et al., 2008). Under the accelerated pace at which the effects of climate change are negatively impacting shallow coral reefs worldwide (França et al., 2020; Pernice and Hughes, 2019), the number of threatened coral species is expected to rise, with rare ones facing increased risk of extinction (Fagan et al., 2005). Species with small geographical ranges, low frequency of occurrence and narrow habitat breadths are more susceptible to both natural and human-induced disturbances (Davies et al., 2004). As local abundance and reproductive success are key in the persistence of these rare species (Vermeij and Grosberg, 2018), knowledge on the status of these traits within subpopulations of the species’ global range becomes essential to plan and implement effective conservation strategies.

The pillar coral *Dendrogyra cylindrus* is a rare reef building species with a high amount of evolutionary history (Curnick et al., 2015). D. *cylindrus* is restricted to the Caribbean basin (Aronson et al., 2008; Finney et al., 2010) and exhibits a combination of life history traits that makes it prone to local extinctions ((Marhaver et al., 2015; Bernal-Sotelo et al., 2019)). Until recently, knowledge on the species was scarce for the entirety of its global range (Aronson et al., 2008). But, the inclusion of the species within the United State’s Endangered Species Act, catalysed conservation actions focused on bridging knowledge gaps on its reproductive ecology and the status of its subpopulation in the country (Fish and Commission, 2013; Marhaver et al., 2015; Neely et al., 2013, 2018; Chan et al., 2019; Neely and Lewis, 2020). This new knowledge depicts a grim reality for the species future. Decline in colony abundance has been estimated above 80% (Kabay, 2016) driving the Florida subpopulation to reproductive extinction (Neely and Lewis, 2020). Moreover, it is hypothesised that the species could be facing a bottleneck due to reproductive failure (Marhaver et al., 2015).

However there still are important knowledge gaps that need be filled to guide effective conservation action for the global species population. For the past fifteen years, the abundance of *Dendrogyra cylindrus* has only been quantitatively assessed in less than 15% of its distribution range (Bruckner and Bruckner, 2006; Clark et al., 2009; Riegl et al., 2009; Rodríguez-Martínez et al., 2010; Neely et al., 2013; Marhaver et al., 2015; Kabay, 2016; Bernal-Sotelo et al., 2019). Most of the status knowledge is limited to the north edge of the species distribution range in Florida (Chan et al., 2019; Kabay, 2016; Neely et al., 2013, 2018; Neely and Lewis, 2020), the Seaflower Biosphere Reserve in Providencia island, off the coast of Nicaragua in the Colombian Caribbean (Acosta and Acevedo, 2006; Bernal-Sotelo et al., 2019) and Curacao (Marhaver et al., 2015; Chan et al., 2019)). Therefore, more information on the status of the species’ national ranges is needed to ensure effective mobilisation of resources at the national and regional level for its conservation.

In Venezuela, data information about the distribution and abundance of the pillar coral was limited to qualitative data gathered in the 1980s (Weil, 2003). Limited to anecdotal accounts from researchers, the species was last assessed as data deficient (DD) on the national Red Book of threatened specie and by 2013, a contraction of at least 50% of its national range was hypothesised. Here, we present the results of a status assessment of *Dendrogyra cylindrus* within the southern edge of the species’ global distribution range, carried out between 2014 and 2016 in Archipelago Los Roques National Park (”Los Roques”) where the species representativeness in relation to the different protection levels of this multi-use MPA, was also evaluated.

## MATERIALS AND METHODS

### Study Area

Los Roques is located north off the Venezuelan coast (N 11 44’26” - 11°58’36”, W 66 32’42” - 66°57’26”, Fig. 1) and was the first MPA in the country, created in 1972 under IUCN category II. The archipelago is semi-enclosed within two barriers that provide a semi-atoll morphology: the eastern (20 km long) and southern (30 km long) barriers. The system comprises 50 coralline cays with fringing and patch reefs, sandbanks, and extensive mangrove forests and seagrass beds (Weil, 2003). The MPA is regarded as one of the most important coral reef systems in the country (Rodríguez-Martínez et al., 2010) and the southern Caribbean (Network, 2014).

**Figure 1.**
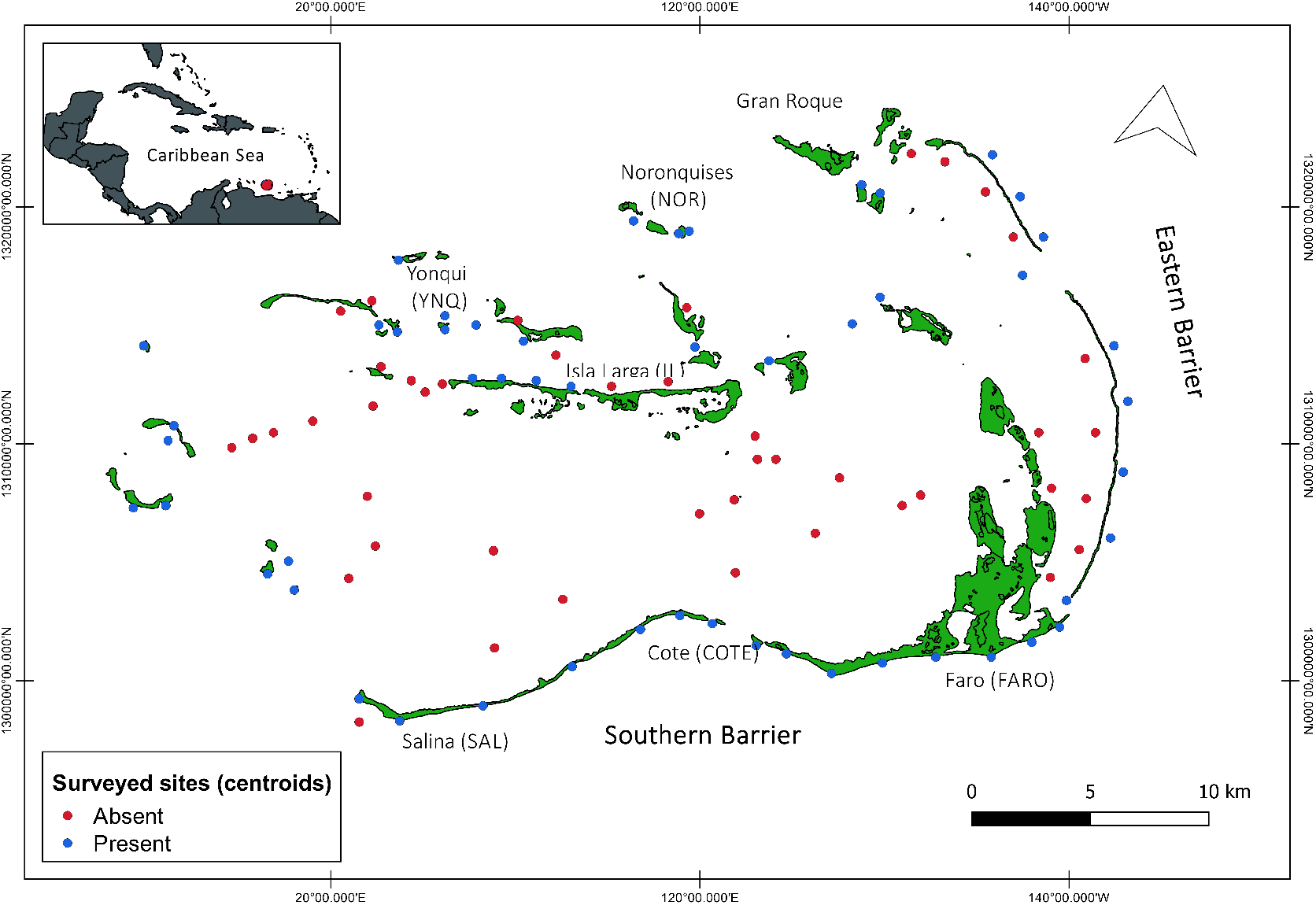
Sites surveyed for the census at Archipelago Los Roques National Park, Southern Caribbean Region. The map extent represents the MPA area of 225.153 ha.

### Distribution and Abundance

To determine the distribution and abundance of *Dendrogyra cylindrus*, we conducted a census across the archipelago between 2014 and 2015, encompassing 106 sites and covering an area of 6.2 Km^2^. These sites included leeward and windward cays, fringing and barrier reefs, reef patches, mixed seagrasses and sand banks, as well as sites indicated by interviewed stakeholders.

We used linear paths covering a 20 m-wide area between 1m and 15m depth at each site. Four freediving observers performed the surveys. Because of the low occurrence of the species, the length of the linear paths varied between 400 and 1,000m according to the extent of the geomorphological formation and/or habitat type. At each path the starting and end points were georeferenced with a Garmin 60S GPS receiver.

The first sites surveyed were identified using local ecological knowledge (LEK) because until 2014, only one colony of the species had been consistently reported for the MPA (Bastidas et al., 2012). For this, questionnaires were administered through a combination of conversational and standardized interviewing (Schnell and Kreuter, 2003) to three different local stakeholders' groups using snowball sampling. These groups included licensed artisanal fishers, SCUBA diving guides from local operators and lodge managers. All lodges, dive operators and fishers congregation points were visited during the study period. We achieved a sampling size of 35%, 100% and 56% out of 200 licensed fishers, three dive operators and 65 lodges, respectively. Based on response time and clarification requests during a pilot study, we included closed and open questions as well as validation ones to reduce social desirability bias (Kreuter et al., 2008). Prior verbal consent by interviewees, the following four questions were asked: 1) Do you know the pillar coral? (yes/no answer), 2) Could you list the numbers that corresponds to the pillar coral? when shown an option card with numbered pictures of local coral species to validate the previous question 3) Have you seen the pillar coral in Los Roques? (yes/no answer), and 4) Where have you seen pillar corals in Los Roques? Open question which included a gridded map of the archipelago where respondents could draw and explain their answers.

From sites identified through LEK, all shallow systems in a 15 Km linear distance from the site were systematically surveyed until all potential occurrence areas for the species within the MPA were surveyed. Within each path, the number of colonies and recently detached fragments, depth, substrate, and slope were annotated. Health status was assessed through partial and total mortality, diseases, and bleaching prevalence based on the presence/absence of visible signs. Every sign of tissue discontinuity (recent or old), disease and/or health problems (i.e. white band, white spot or patchy necrosis) was registered using ID cards (Weil and Hooten, 2008). Colony abundance was assessed on the count of discrete units taking no tissue and/or skeleton connection as the criterion to distinguish among units. Additionally, wave exposure (windward and leeward) and geomorphological unit (i.e. fore reef, back reef, reef terrace, and lagoon) were recorded at each site.

### Habitat Characterization

Habitat characterization at each site was assessed qualitatively according to the habitat type, habitat condition and geomorphological unit. Habitat type was based on the main substrate and dominance of benthic species following Croquer et al. (2016) and Bernal-Sotelo et al. (2019). This rapid characterization is suitable for rough descriptions of the coral benthic communities at spatial scales above 50 m (Hill and Wilkinson, 2004) as in this study. According to the relative predominance of live coral cover, macro-algae, bare substratum and sedimentation, each site was assigned to one of four habitat condition categories. These included: (1) ‘Excellent’ (i.e. the corals clearly dominated the benthos, with no evidence of sedimentation and/or coral mortality), (2) ‘Good’ (corals still dominated, sedimentation was evident but dead corals were rare), (3) ‘Regular’ (corals and fleshy macroalgae have similar abundance and sedimentation was obvious, bared substratum was common) and (4) ‘Degraded’ (fleshy macroalgae clearly dominated the benthos with sediment often smothering corals and bare substratum was abundant). Seascape complexity was visually estimated on a scale of 1 (flat, little relief) to 5 (highly complex) following Polunin and Roberts (1993). Post-survey standardization based on the range of seascape complexity was performed based on photographs for all surveyed sites (Newman et al., 2015) and 3D models at reef scales from six sites (Agudo et al., 2019). This method was employed as it has been widely used (Polunin and Roberts, 1993; Jennings et al., 1996; Wilson et al., 2007; Newman et al., 2015) and allows to assess complexity in a large-scale survey as in this study. Prior to the census, observers calibrated the application of qualitative assessment criteria for habitat until no difference among them was obtained.

### Size Structure

Size structure of *Dendrogyra cylindrus* population(s) in Los Roques was determined by measuring maximum height of each colony within 20 sites where *D. cylindrus* occurred across the archipelago following Acosta and Acevedo (2006). Additionally, demographic disruptions as described in Sodhi and Ehrlich (2010) were investigated at spatial scales of thousand and hundreds of meters according to habitat type to assess differences in population dynamics at these scales. Although size structure represents a stage in a dynamic process of population growth and decline (Bak and Meesters, 1998, 1999; Crabbe, 2009), it can be used to identify conservation or management units in the absence of more detailed population biology data and restrained resources to obtain them. This is because corals’ size structure carries demographic information (Hughes and Connell, 1987; Elahi and Edmunds, 2007; Crabbe, 2009) and can reflect differences in natural and anthropogenic stressors at the reef scale (Bak and Meesters, 1998, 1999).

To assess demographic disruptions among sites and habitat types, a hierarchical design with two factors was employed considering habitat as a fixed factor and reefs nested within habitat as a random factor. The two most common habitat types where the species was found were used as levels [crossed with two levels: Sand Flats (SF) and Consolidated Reefs (CR), separated by 10s km], randomly selecting three sites for each of those habitats as levels of the factor reef [random with six levels: Noronqui (NOR), Yonqui (YNQ) and Isla Larga (IL) within SF, and Salina (SAL), Cote (COTE) and Faro (FARO) within CR, separated by 100s m]. Between April and November 2015, four 50m x 10m belt transects were placed parallel to the coast on the 6m-8m depth contour at each reef, totalling a surveyed area of 2,000*m*^2^. Transects were systematically laid from a randomly chosen point, following the predominant current. Colony maximum height was measured with a tape (±0.1*cm*). Overturned colonies and fragmented pillars that resumed vertical growth (Fig. 2a) were also recorded. As fragments and overturned colonies have lower reproductive output -if at all- (Lirman, 2000), these were excluded from the analyses to reduce biases on estimating subpopulation size. Here we use the term subpopulation and global population following the IUCN Red List of Threatened Species Guidelines version 14 (https://www.iucnredlist.org/resources/redlistguidelines).

**Figure 2.**
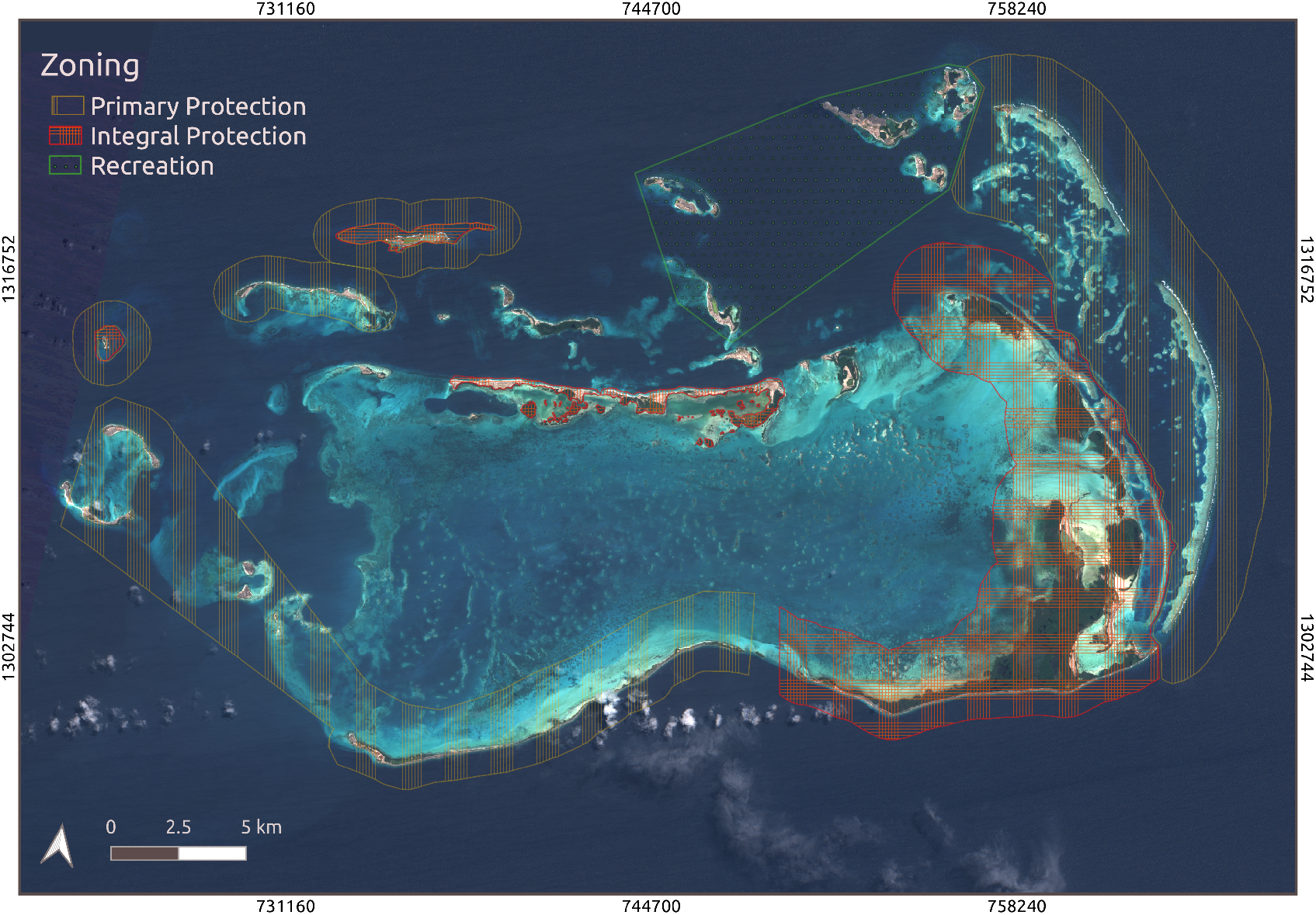
Archipelago Los Roques National park zoning, according to use regulations established in 1991. Based on allowed activities zones form higher to lower protection are: Integral Protection (IPZ), Primary Protection (PPZ), Marine Manage Area (MMA - corresponding to the extent not included in the other zones) and Recreation (ReZ). Image contains modified Copernicus Sentinel data [2020], processed by ESA.

Size frequency distributions were built for the MPA and for each reef based on colony maximum height. Because population dynamics (Hughes, 1984; Edmunds, 2002; Elahi and Edmunds, 2007) and intrinsic factors affecting growth(Hughes and Connell, 1987) were not assessed, we chose systematic size classes only at the MPA scale based on those used by Acosta and Acevedo (2006). At the other six sites (reef scale) asymmetrical size classes were used to maximize the number of colonies per class. For the latter, seven size classes were established as follows: class I, composed by juveniles (colonies < 15*cm* in maximum height and isolated from any other colony; (Acosta and Acevedo, 2006), small colonies encompassing class II (16 – 25*cm*), class III (26 – 50*cm*) and IV (51 – 100*cm*), and medium and large colonies defined as 50% of the height range determined in the study and encompassing classes V (101 – 150*cm*), VI (151 – 200*cm*) and VII (> 200*cm*).

### Representativeness within the MPA

To evaluate *Dendrogyra cylindrus’* representativeness in Los Roques *sensu* Margules and Pressey (2000), a gap analysis was performed utilising vector layers of the MPA zoning, the species area of occurrence and the habitat types and condition described for all surveyed sites. The MPA zoning layer was built following the National Park’s use regulation plan (PORU by its Spanish acronym; Official Gazette No. 4250, 1991) where each of the different protection levels by zone are defined and delimited. According to the PORU, except for Gran Roque island, the MPA’s marine systems’ protection level falls within one of five zones (Fig. 2). 1) the Integral Protection Zone (IPZ) represents the highest level of protection; only scientific research and environmental protection activities are allowed. 2) the Primary Protection Zone (PPZ) where nature interpretation for educational purposes and nature-based tourism activities are allowed with a maximum group size of 15 people and previous authorisation. 3) the Marine Managed Area encompassing the biggest area within the MPA where passive recreation activities and small-scale fishing are allowed without the use of nets or cages. 4) the Recreation Zone limited to the north-east cays and islands of the archipelago; here infrastructure to service tourism and recreation is permitted and activities such as sport fishing, water sports and the free transit of boats take place.5) The navigation channel which is a special-use area designating the route that need be followed by vessels and boats entering the archipelago from the south. The GIS desktop software QGIS 2.4 Chugiak and the package “sp” (Pebesma et al., 2020) implemented in R (Team, 2017) were used to built all vector layers and their intersections.

To assess the spatial structure of the species representativeness, habitat complementarity or redundancy, and habitat diversity within each zone of the MPA were estimated at a landscape scale, while percentage landscape occupancy and nearest neighbour distances were estimated using all habitat polygons as patches. For this the software FRAGSTAT (McGarigal et al 2012) and the package “raster” (Hijmans et al., 2020) were used. Layers’ extent was set to 221,120 hectares, corresponding to the MPA area and using a 200 m^2^ grain (56% of the area from the smallest habitat polygon) at a scale of 1: 200,000. Following Halpern et al. (2015), a modified Sorensen similarity index was used to estimate habitat complementarity.

### Data Analysis

#### Distribution and Abundance

A distribution map for *Dendrogyra cylindrus* at the MPA scale was produced. Surveyed area at each site was obtained through geometry tools from the construction of polygons based on the georeferenced belt transects in a vector layer. For each site, the number of colonies, colony density, partial mortality, bleaching, and disease prevalence as well as habitat type, habitat health status and complexity were used as attributes. Colony density per site was estimated by standardizing the number of colonies by polygon area. Relative abundance of *D. cylindrus* colonies per category of all the habitat attributes assessed in this study was standardized by the number of sites assigned to each category.

To assess the relative importance of wave exposure, reef location, depth, slope, substrate, habitat type, habitat condition and structural complexity on the abundance and occurrence of *D. cylindrus* at the MPA scale, gradient boosted regression trees (GBRT) were used. This technique can handle many types of response (numerical, categorical, censored and dissimilarity matrices) variables and model complex interactions among predictors as well as non-linear relationships (De’ath, 2007), proving a useful analytical tool to identify relevant explanatory variables for observed patterns in ecological studies (Elith et al., 2008; Maloney et al., 2012). Two GBRTs were fitted predicting the abundance and occurrence of *D. cylindrus colonies*, with colony density per site and presence/absence data as response variables, respectively. For this the “gbm” package (Greenwell et al., 2020) and functions written by Elith et al. (2008) were used. The H2O open source AI platform (v 3.10.1.2, www.h2o.ai) implemented in R (Team, 2017) was used for parameter tuning. Habitat types that were assigned to less than five sites as well as those sites with less than three colonies were dropped during modelling to avoid modelled relationships based on sparse data.

Model performance was based on the following metrics: area under the receiver operator characteristic curve (AUC) (Rosset, 2004a) for *Dendrogyra cylindrus* occurrence and the Akaike Information Criteria (AIC) and deviance for colony abundance. AUC estimate how well-fitted values discriminate between observed presences and absences, with values ranging from 0.5 (no better than random) to 1.0 (perfect discrimination). Model comparison and parameter tuning (number of trees, tree depth, learning rate, and bag fraction) were performed through a random grid search with a 10-fold cross validation on randomly withheld data using convergence-based early stopping for performance metrics (rounds: 3, tolerance: 0.01)(Aiello et al., 2015). The models that showed highest average scores with lowest standard deviation values on performance measures from the cross validation were later evaluated. Here, models with AUC scores > 0.8 and > 0.9 are considered very good and excellent, respectively (Rosset, 2004b). The relative importance of each predictor variable was estimated based on the number of times a variable was selected for splits and weighted by the squared improvement of the model, averaged over all trees where higher numbers indicate a stronger influence on the response variable (Friedman and Meulman, 2003). Relative importance values were scaled to 100.

#### Size Structure

To test the null hypothesis of no demographic disruptions at the two spatial scales evaluated, a permutational multivariate analysis of variance (PERMANOVA) was performed from dissimilarity matrices based on a distribution overlap index (DO) following Ernest (2005). For this, size class frequencies were treated as variables. Non-metric multidimensional scaling (nMDS) ordinations were used to illustrate dissimilarities on size frequency distributions among levels of the factors examined. These analyses were performed using PRIMER + PERMANOVA v6 software (Clarke and Gorley, 2006) while the dissimilarity matrices were constructed using the “vegan” package (Oksanen et al., 2019). A SIMPER test was also performed to identify the size classes that contributed the most to dissimilarities between size frequency distributions.

#### Representativeness

Logistic regressions using a cumulative model were performed to test whether the MPA zoning had an effect on the abundance of *Dendrogyra cylindrus* colonies, partial mortality prevalence and habitat condition. For the latter non-standardised cut points were used for the four habitat condition categories (Agresti, 2010). The null hypothesis of no difference in (1) colony abundance, (2) partial mortality prevalence and (3) habitat condition among use zones was tested using a chi-square test. For this, abundance and partial mortality were transformed to binary variables, where a site was assigned a value of 1 if the number of colonies exceeded or equaled that of the MPA’s average and the frequency of partial mortality was below 30% following the criteria used to categorise habitat condition. Sites where none of these conditions were met, were assigned a value of 0. All analyses were performed using packages “caTools” (Tuszynski, 2020) and “ordinal” (Christensen, 2019).

## RESULTS

### Distribution and Abundance

A total of 6.2km^2^ were surveyed encompassing developed, marginal, barrier and patch reefs as well as sand flats and seagrass patches. Within 49% out of the 106 surveyed sites 1,490 discrete units (colonies) of *Dendrogyra cylindrus* were counted in 719 sightings. Colony clusters varied from small five-colonies aggregations to two isolated clusters of more than 100 colonies on sand flats in the central north cays of the archipelago, forming mono-specific reefs exceeding 100 m^2^ in area (Fig. 3b). Seventy percent of surveyed sites were identified through LEK. Only 6.3% of fishers’ responses did not coincide with the observed species’ occurrence. Moreover, 30% of interviewed fishers further elaborated on their knowledge, relating *D. cylindrus* colonies with spiny lobster fishing sites: *”we often find the lobsters between the base of the caramujo (local name for the species) and the sand. Our best fishing sites are those with a lot of caramujos”* acknowledging the species importance for the most profitable local fishery.

**Figure 3.**
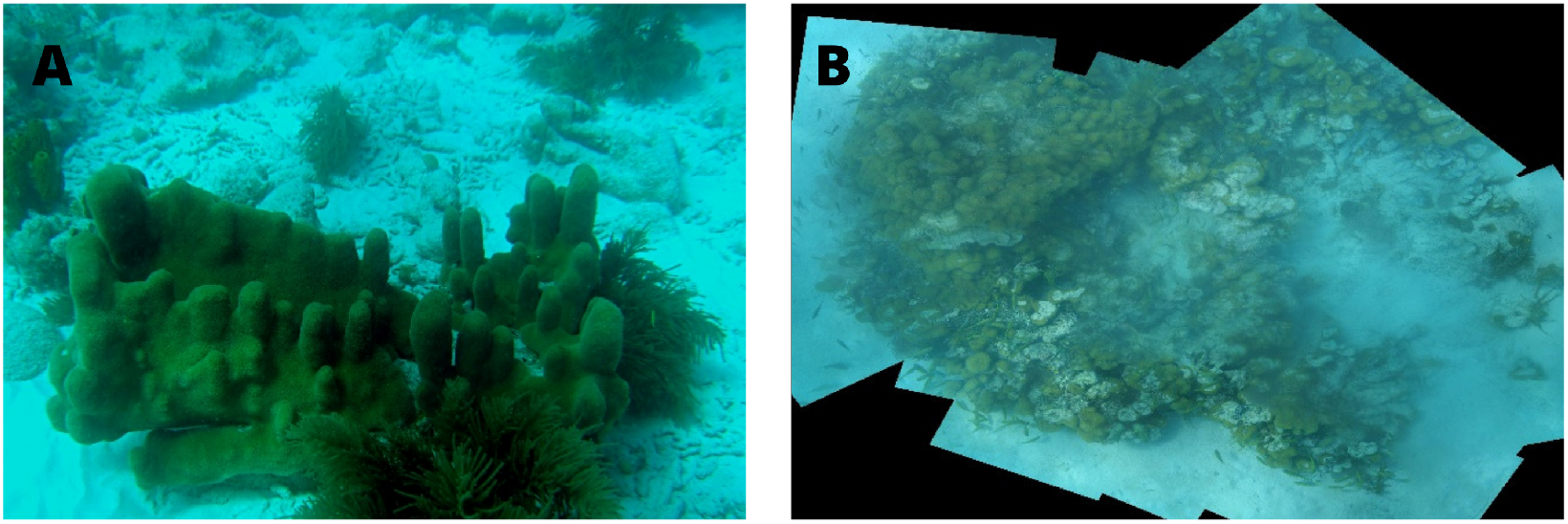
Overturned colony of *Dendrogyra cylindrus* with resumed vertical growth (a); Photomosaic of a monospecific reef of *D. cylindrus* in a sand flat near Espenqui at Los Roques (b).

Knowledge on the species varied according to the respondent’s group. While 79% of the interviewed fishers correctly identified the pillar coral, only 12% and 13% of lodge managers and dive guides, respectively, were able to do so. Fishers who identified the pillar coral engaged in spiny lobster fishing on a yearly basis. All SCUBA diving guides recognized the pillar coral colony from the numbered photographs card and reported to have seen it regularly in two of the most frequented dive sites of Los Roques. One of those sites, Boca de Cote had the second highest colony abundance in our surveys (Fig. 4). Only 5% of respondents failed the validation question. Colony abundance ranged widely between 1 and 68 colonies per site, with an average of 16.3 ± 35.23 (mean ± SD). Highest colony abundance was observed at three sites in the southern barrier (177, 151 and 154, respectively). However, when standardized by surveyed area, consolidated reef sites in the southern barrier showed lower densities (0.63/ 100*m*^2^ ± 0.5, *mean* ± *SD*) than sand flat sites in the central northern cays (5/ 100*m*^2^ ± 3, *mean* ± *SD*, Fig. 3).

**Figure 4.**
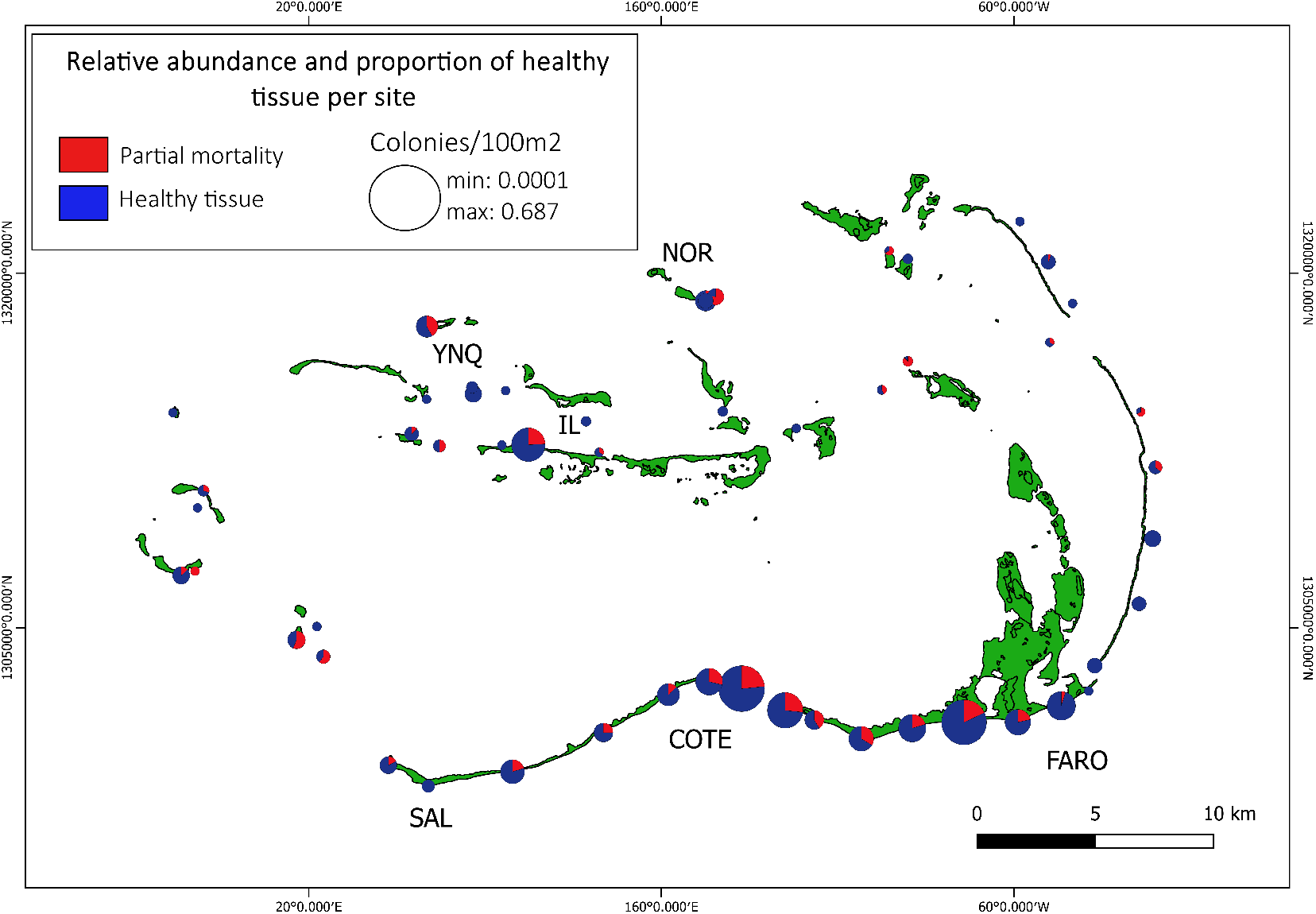
Proportion of *Dendrogyra cylindrus* colonies with partial mortality based on the relative density (colonies/100m^2^; area of pie charts) of colonies observed at 56 out of 106 sites across Los Roques. Noronqui (NOR), Yonqui (YNQ) and Isla Larga (IL) correspond to Sand Flat sites and Salina (SAL), Boca de Cote (COTE) and El Faro (FARO) to consolidated reefs sites

Generally, once we found a colony, there were more colonies in less than a 5 m radius. The mean number of colonies per cluster was 2.83 ± 1.63. This might reflect either a close-to-adult recruitment, a high rate of asexual reproduction by fragmentation, or both. Nevertheless, “juvenile” (< 15*cm* height)(Riegl et al., 2009) and big colonies (> 150*cm* height) were never observed as part of clusters, suggesting a high rate of fragmentation among medium sized colonies. In total, 15 juvenile colonies were counted, 200 recently detached pillars or fragments, and 15 recognizable overturned colonies were observed across the MPA. Overall, syndromes and disease prevalence as well as partial mortality were low on *Dendrogyra cylindrus* across the MPA. Out of the 1,490 colonies observed, only 28.9% showed partial mortality, whereas 0.2% and 0.27% showed signs of white plague and black band disease, respectively. A total of five colonies with bleaching signs across all sites, and six colonies with total mortality -all > 100*cm* height-in sites within the Southern Barrier were counted.

### Habitat

A total of 14 different habitats were described across all surveyed sites (SM Table 1). *Dendrogyra cylindrus* occurred in all described habitats except for seagrass patches and sand flats dominated by “weedy” corals (i.e. *Millepora* sp. and *Porites* sp). GBRT models indicated that habitat characteristics had the strongest influence on both occurrence and abundance (Table 1). Relative importance of habitat condition and type of habitat accounted for 60% of the variability in the probability of occurrence and the abundance, however for the latter, the relative importance of geomorphological units surpass that 8/20 of habitat condition, which is expected as this variable is determinant at settlement. Most interactions among predictive variables were very low and non-significant, with the exception of type and condition of habitat (H=0.34, p=0.03) for the occurrence model, and habitat condition with both, complexity and geomorphological unit (H=0.56, p=0.04 and H=0.25, p=0.05, respectively) for the abundance model. Model performance was better for occurrence (AUC=0.94) than for abundance (AIC=0.89).

**Table 1.** Relative importance of habitat variables used to explain *Dendrogyra cylindrus* occurrence and colony abundance at Archipelago Los Roques National Park, using adjusted Gradient Boosted Regression Trees. Values are expressed in percentage (%)

| Variable | Abundance | Occurrence |
| --- | --- | --- |
| Habitat health condition | 33.10 | 33.43 |
| Habitat type | 26.39 | 17.04 |
| Geomorphological unit | 23.42 | 22.16 |
| Qualitative structural complexity | 5.47 | 5.78 |
| Depth | 4.23 | 4.57 |
| Slope | 4.01 | 3.95 |
| Substrate | 3.25 | 12.85 |

The fitted functions revealed that the occurrence probability was greater on consolidated reefs dominated by *Orbicella* sp. or *Pseudoplexaura* sp. (97.5%) and sand flats either dominated by *Pseudoplexaura* sp. or *Acropora cervicornis* (95.3%) in Excellent (98%) and Good (85%) condition. Higher abundances were observed in complex consolidated reefs dominated by *Orbicella annularis* (26.51%). This type of habitat prevailed in the southern barrier topography. Abundance was also high in moderately complex sand flats dominated by octocorals (17.56%) where *Dendrogyra cylindrus* was the species contributing the most to structural complexity, as well as in patches of *Acropora cervicornis* (13.91%).

Regarding habitat condition, most sites where *Dendrogyra cylindrus* was observed, were classified as Good or Excellent (79.69% and 10.57%, respectively). Only 38% of all habitats with or without *D. cylindrus* were classified within the Regular and Degraded categories. Similarly, the abundance of colonies, normalized by the number of sites in each category, was higher on the Good and Excellent habitat categories, while partial mortality prevalence, also normalized by the number of colonies per site, was higher on the Regular and Degraded habitat categories. Contrasting with previous studies (Acosta and Acevedo, 2006), the number of colonies was five times greater on windward (83.9%) than leeward (16.10%) sites. Frequency of observed colonies was higher at a 4.5m-8m depth interval. Similarly, the frequency of occurrence was higher in reef terraces (62.45%), followed by front (13.35%) and back (10.85%) reefs, whereas no colonies were observed inside the lagoon.

### Size Structure

Colony height varied between 5cm and 290cm, with and average of 71.6 ± 93.84 (Table S2). Size frequency distribution for the MPA was symmetric (mode= 32 cm), though most of the colonies were below 60 cm height (Fig. 4). Average colony height differed between habitat types (Pseudo-F=2.95, p=0.048, Fig. 4), with consolidated reefs showing taller colonies than sand flats (Fig. 5). However, size structure differed only at the reef scale (Table 2), with some reefs like El Faro having mostly taller colonies (median= 126.5cm; mode=260cm) and Yonqui, having mostly small and medium ones (median= 31cm; mode=63cm; Table S2). These results suggest that the growth rate might be dependent on factors relating to the type of habitat and/or operating at spatial scales larger than reefs, whilst population dynamics such as mortality and recruitment (accounting also for fragments) might be determined by factors operating at reefs scales. SIMPER analyses showed that these differences were due to the presence of smaller size classes (most recently recruited colonies) in few consolidated reefs and a low frequency of colonies within medium size classes in El Faro. In contrast, most sand flat reefs lacked colonies above 100cm, being dominated by colonies within the small and medium size classes (Fig. 5).

**Figure 5.**
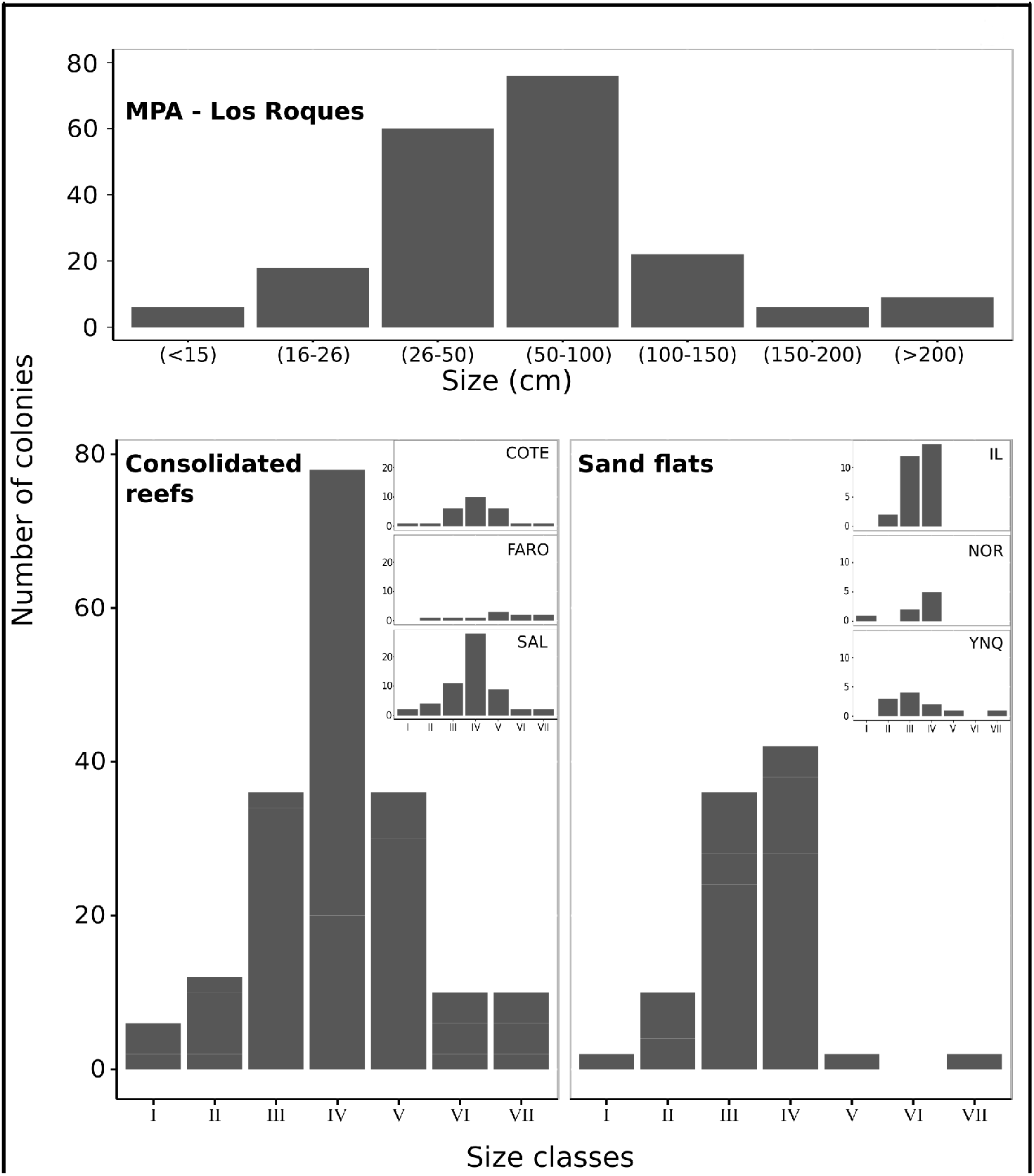
Colony size (maximum height) distribution of *Dendrogyra cylindrus* in Los Roques Archipelago National Park and six sites, three at each of the two most common habitat types for the species at the MPA: consolidated reefs (Boca de Cote:COTE; Salina: SAL; and El Faro: FARO) and sand flats (Isla Larga: IL; Noronqui: NOR; and Yonqui: YNQ).

**Table 2.** Analysis of Variance based on permutations (PERMANOVA) of similarity matrices of Ernest (2005) overlapping index calculated using the frequency of *Dendrogyra cylindrus* colonies within each of seven size classes in six sites (nested factor) within consolidated reefs and sand flats (fixed factor habitat) at Archipelago Los Roques National Park.

| Source of variation | df | Mean Square | Pseudo-F | P-perm |
| --- | --- | --- | --- | --- |
| Habitat | 1 | 161.33 | 1.6916 | 0.304 |
| Reef(Habitat) | 4 | 95.375 | 2.7057 | 0.036 |
| Residuals | 18 | 35.25 |  |  |
| Total | 23 |  |  |  |

### Representativeness

Most of the surveyed habitat’s area in Los Roques lies inside the MMA (55.50%) and Recreational (21.30%) zones where activities susceptible to be localised threats (Aronson et al., 2008; Mumby et al., 2014) are carried out. This is also true for some key habitats such as those dominated by *Acropora cervicornis*, where over 70% of its area is inside the MMA (52.6%) and Recreational (47.29%) zones. *Dendrogyra cylindrus’* area of occupancy represent 1.07% that of the MPA and is somewhat evenly distributed among the non-take zones or IPZ (24.4%) and PPZ (31.18%) and the lower protection level MMA zone (31.18%, Fig. 6). This was also true for colony abundance, with 51.48% of colonies inside the MMA zone and 43.48% inside the non-take ones (23.59% and 19.79% for IPZ and PPZ, respectively).

**Figure 6.**
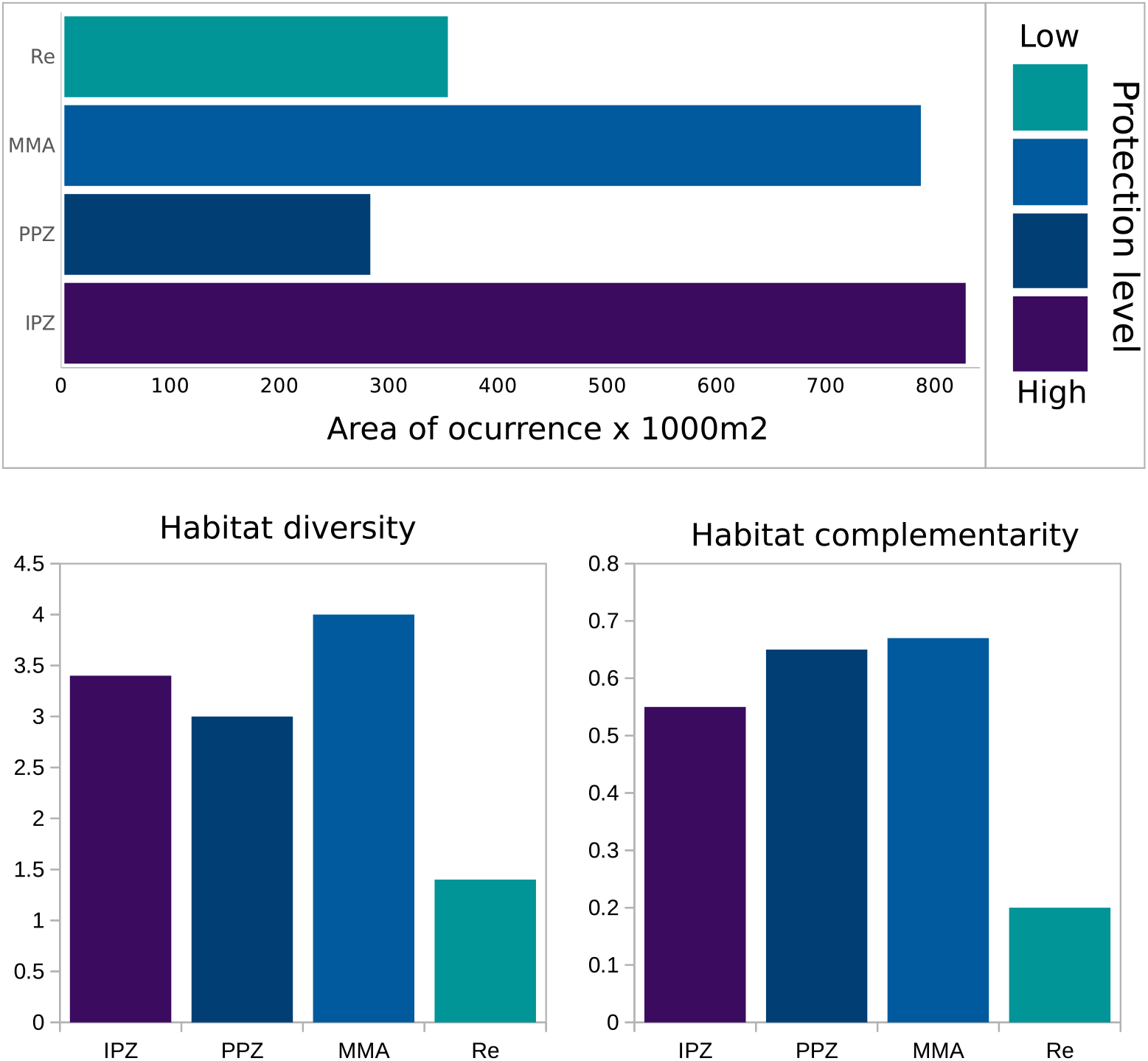
Area of occurrence, diversity and complemetarity of *Dendrogyra cylindrus* habitat according to the zoning regulating uses and activities in Archipelago Los Roques National Park.

The MMA zone had the highest value of habitat diversity (*H* = 3.949), followed by the non-take IPZ and PPZ zones (*H* = 3.323 and *H* = 2.990, respectively). Only the IPZ encompass all six habitats where *Dendrogyra cylindrus* abundance surpassed 16.30± 35.23 (*mean* ± SD) colonies per site, however, habitat complementarity or redundancy was higher within the MMA (C=0.67) and the PP (C= 0.65) zones (Fig. 6). Only four occurrence sites are below an average distance of 100 m from other sites with the same type of habitat; all consolidated reefs and pavement terraces in the southern and eastern barriers, respectively. Sand flat occurrence sites had the greatest average distance to their five nearest neighbours (1.45 ± 0.87*km*; *mean* ± SD). MPA zoning was a good predictor of colony abundance (*X*^2^ = 10.20; *p =* 0.037) and habitat condition (*X*^2^ = 27.17; *p =* 0.007); however, model sensitivity for colony abundance was low (25%) and only the Recreational zone had a significant effect over the condition of habitats (Table 3).

**Table 3.**
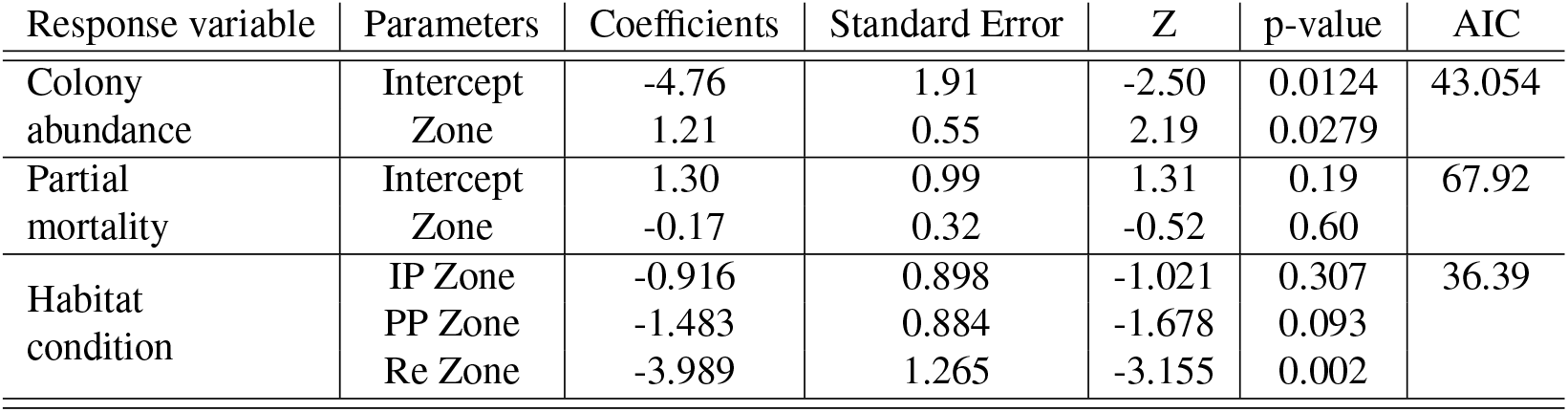
Model parameters describing the effect of the zoning regulating uses and activities at Archipelago Los Roques National Park on the abundance of colonies, partial mortality prevalence and habitat condition of *Dendrogyra cylindrus*.

| Response variable | Parameters | Coefficients | Standard Error | Z | p-value | AIC |
| --- | --- | --- | --- | --- | --- | --- |
| Colony abundance | Intercept | -4.76 | 1.91 | -2.50 | 0.0124 | 43.054 |
|  | Zone | 1.21 | 0.55 | 2.19 | 0.0279 |  |
| Partial mortality | Intercept | 1.30 | 0.99 | 1.31 | 0.19 | 67.92 |
|  | Zone | -0.17 | 0.32 | -0.52 | 0.60 |  |
| Habitat condition | IP Zone | -0.916 | 0.898 | -1.021 | 0.307 | 36.39 |
|  | PP Zone | -1.483 | 0.884 | -1.678 | 0.093 |  |
|  | Re Zone | -3.989 | 1.265 | -3.155 | 0.002 |  |

## DISCUSSION

We assessed the status of *Dendrogyra cylindrus* in the southernmost portion of its distribution range for the first time. We reported over 1,000 colonies in 14 different habitats where the species is conspicuous, but infrequent. Taking into account colony abundance, health status and size structure, our results suggest that Los Roques could be a stronghold for the global population of *D. cylindrus*.

The number of colonies recorded in this study is the highest reported for the species to date (Szmant, 1986; Karpouzli et al., 2004; Quinn and Kojis, 2005; Acosta and Acevedo, 2006; Clark et al., 2009; Riegl et al., 2009; Rodríguez-Martínez et al., 2010; Neely et al., 2013, 2018; Neely and Lewis, 2020; Marhaver et al., 2015; Chan et al., 2019; Galford et al., 2018; Bernal-Sotelo et al., 2019), even when compared to studies where large-scale assessment to identify occurrence sites have been done (Acosta and Acevedo, 2006; Bernal-Sotelo et al., 2019). However, this difference could probably respond to the larger sampling effort made in Los Roques. Similarly, colony size structure for the whole MPA suggests *Dendrogyra cylindrus* could maintain a positive population growth rate (Riegl et al., 2003) at Los Roques in the absence of a catastrophic event. The unimodality of the size distribution at the MPA scale suggests this subpopulation has not been impacted by episodic disturbances (Edmunds and Elahi, 2007) while the dominance of small and medium size colonies (26 100*cm* height) suggest a high fecundity potential (Babcock, 1991; Nozawa and Lin, 2014) with lower incidence of partial mortality; conferring potential for intrinsic population growth (Bythell et al., 1993, 2018; Edmunds and Elahi, 2007).

However, as *Dendrogyra cylindrus* size structure showed differences among reefs, our results also suggest that dynamics among reefs are asynchronous and thus, intrinsic population growth for the MPA is dependent upon reef-level processes affecting population parameters at occurrence sites. Under low sexual recruitment as it has been observed for the species (Marhaver et al., 2015; Neely and Lewis, 2020) this makes Los Roques subpopulation highly vulnerable to site-level or local extinctions; something that has been previously observed in other MPAs. In the SBR, between 2002 and 2012 *Dendrogyra cylindrus* suffered local extinction in 50% of the plots surveyed in both years, albeit increasing the total number of colonies and fragments (Bernal-Sotelo et al., 2019).

Contrary to most consolidated reefs sites, dominance of small and medium colonies in sand flat habitats within the archipelago evidence periods of no transition among size classes or of high clone propagation (Lirman, 2003; Riegl et al., 2003; Chan et al., 2019; Bernal-Sotelo et al., 2019), suggesting these sites might have a higher number of clones. High clonality in *Dendrogyra cylindrus* subpopulations has led to reproductive extinction in Florida (Neely and Lewis, 2020) and is thought to be one of the main factors leading the species to a bottleneck (Marhaver et al., 2015). However, these sites could be important for the species persistence within the MPA. Fragmentation also allows to maintain positive intrinsic rates of population growth in Scleractinia species with low recruitment rates (Lirman, 2000, 2003; Edmunds and Elahi, 2007; Edmunds, 2015; Foster et al., 2013) and facilitates increasing the extension of occupancy (Lande et al., 2003; Reed, 2005) either through a higher survival of fragment (Lirman, 2000) or by providing adequate substrate for sexually produced larvae to settle (Vermeij, 2005; Ricardo et al., 2017). The latter is especially true for sand flat habitats such as Noronqui and Isla Larga, where a high number of colonies were found. Though, it is necessary to assess the proportion of clones per site as well as the degree of connectivity among them, the average distance between the first five neighbour occurrence sites in sand flat habitats, exceeded that reported as maximum dispersion distance for clones in Florida (Chan et al., 2019). Additionally, the presence of small juvenile colonies in Los Roques suggests that some degree of sexual recruitment has occurred in the recent past.

Differences found between habitat types, habitat condition categories and at reef scales in both abundance and population structure have important implications for management. This is because achieving representativeness for the species within high protection zones entails accounting for these differences as well as the varying degree of exposure and susceptibility to both localised and global threats within occurrence sites.

Our results suggest that zoning in Los Roques has been effective in maintaining the MPA’s conservation value for coral reefs, also providing representativeness to *Dendrogyra cylindrus*. However, the area of occupancy, habitat complementarity or redundancy and habitat singularity need be accounted too when evaluating how the configuration of the species representativeness might affect its persistence. For example, occupancy of a wide variety of habitats and the growth plasticity observed in *Dendrogyra cylindrus* at Los Roques, could be a testament to the species ability to colonise various habitats (Rogers, 2017; Bernal-Sotelo et al., 2019). But the relatively low number of colonies observed in most of these habitats suggests that dispersion ability does not necessarily translate into population growth or persistence. The number of colonies of the species in habitats other than consolidated reefs and sand flats dominated by octocorals-antipatharia rarely surpassed 10.

Indeed, *Dendrogyra cylindrus* status is strongly determined by the type and quality of habitat, at least in Los Roques. Results from the GBRT indicated that the habitat’s health status and type were the most important variables determining both occurrence and abundance. This makes *D. cylindrus*’ persistence highly vulnerable to habitat degradation and reinforces the importance of site-level management to reduce local or site-level extinctions and mitigate their impacts in the MPA subpopulation. The majority of the 1.07% MPA area occupied by the species is within the zones with lowest protection, including singular habitats in excellent condition such as thickets of *Acropora cervicornis* in the Recreation zone and a mound of *Madracis* sp. in the Navigation channel. These low-protection zones also had the highest redundancy of habitats. Therefore, to effectively achieve representativeness of *Dendrogyra cylindrus* and reef habitats in a manner that increases the chances of persistence for the species, the Integral Protection zones should be extended to contain a habitat complementarity above 0.7 for the four main habitats of the species as well as the singular habitats recorded here, which have disappeared in most of the Caribbean.

Accounting for the prevalence of diseases, bleaching and partial mortality, colony abundance and size structure, the status of *Dendrogyra cylindrus* subpopulation in Los Roques seems better than other areas of its global range (Acosta and Acevedo, 2006; Alvarez-Filip et al., 2019; Neely et al., 2013; Neely and Lewis, 2020; Bernal-Sotelo et al., 2019). However, we cannot conclude in terms of effective population size, for the number of genets and other traits relevant to the species viability (Marhaver et al., 2015; Neely and Lewis, 2020) were not assessed here.

Los Roques has been suggested as a stronghold for other shallow reef building coral species in the Caribbean due to the combination of: 1) a historic low frequency and extent of episodic disturbances, 2) no-take zones established over 30 years ago and 3) good enforcement (Zubillaga et al., 2008; Debrot et al., 2007; Network, 2014). Sadly, in 2011 a drastic change in governance decreased the enforcement capacity of the management authority in over 90% compared to that of the previous five years (National park Superintendent comm. pers. 2015). This is significantly undermining the conservation value of Los Roques and imposing a sense of urgency into investigating the role their shallow reefs-building subpopulations might have for the persistence of other threatened coral species in the southern Caribbean either because they can provide sexually produced larvae or genets for ex-situ conservation interventions.

Here we also prove the value that local ecological knowledge (LEK) has on helping fill knowledge gaps on rare coral species. LEK has been a valuable tool to identify temporal and spatial variations in small-scale fisheries and resource use (GARCÍA-QUIJANO, 2007; Noble et al., 2020) as well as to help update the status and distribution of rare species; being one of the first steps to confirm the presence of species thought to be extinct (Bonfil et al., 2018). Harnessing this knowledge in Los Roques allowed us to confirm the presence of the species in the archipelago, planning surveys in a more cost/effective fashion. Also we were able to record important and new information about the species cultural value among spiny lobster fishers; which could help facilitate participation of fishers in future conservation initiatives.

## ACKNOWLEDGEMENTS

This project was partially funded by an EDGE of Existence Fellowship awarded to Francoise Cavada-Blanco in 2013 and by the Laboratorio de Ecologia Experimental, Departamento de Estudios Ambientales, Universidad Simon Bolivar, through in-kind contributions. The funders had no role in study design, data collection and analysis, decision to publish, or preparation of the manuscript. Field studies were authorized by the Ministerio del poder popular de Ecosocialismo, Hábitat y Vivienda (Approval number: 0323) and Territorio Insular Miranda (Approval number 006). We would like to acknowledge the information provided by local MPA authorities and staff as well as fishers, dive operators and lodges at the time of the study. Their good disposition and courage was key to project completion.

---

**Table S1.** Qualitative characterisation of the different habitat types where *Dendrogyra cylindrus* was most frequently sighted across Archipelago Los Roques National Park between April 2014 and February 2016.

| Substrate | Dominant benthic species | Description |
| --- | --- | --- |
| Inter-reef sand flats | Acropora cervicornis | Low slopes with sparse massive colonies of *Pseudodiploria labyrinthiformis*, *P. clivosa* and *P. strigosa*, *Colpophyllia natans*, *Orbicella annularis* and *Porites astreoides*. Scarce soft corals such as *Pseudopterogorgia* sp. and *Gorgonia ventalina*. Low abundance of carbonate macroalgae of the genus *Halimeda* and *Turbinaria*. |
|  | Disperse massive stony coral colonies | Low slopes with big (1 – 3m height) sparse colonies of *Pseudodiploria labyrinthiformis*, *P. clivosa* and *P. strigosa*, *Colpophyllia natans*, *Orbicella annularis* and *Porites astreoides*, *Acropora cervicornis* and *A. palmata*. Low abundance of carbonate macroalgae of the genus *Halimeda* and *Turbinaria* |
|  | “Weedy” corals | dominated by corals of the genus *Millepora* sp. and *Porites* sp. with *Halimeda* sp. and other fleshy macroalgae such as *Padina* sp. |
| Dead *Acropora palmata* | *Orbicella annularis* | Almost fused fragments of *Acropora palmata* with a mild slope with colonies of *O. faveolata* and sparse massive colonies of *Pseudodiploria labyrinthiformis*, *P. clivosa* and *P. strigosa*, and *Colpophyllia natans*. Scarce soft corals such as *Pseudopterogorgia* sp. and *Gorgonia ventalina*. Abundant carbonate macroalgae of the genus *Halimeda*. |
|  | Storm terraces | Fused fragments of *Acropora palmata*. Steep slope on windward of cays with small colonies (< 50cm height) of *Pseudodiploria labyrinthiformis*, *P. clivosa* and *P. strigosa*, *Colpophyllia natans* and *Porites astreoides*. Abundant carbonate macroalgae of the genus *Halimeda*. |
| Dead Consolidated reefs | Dead *Acropora palmata* | Fused fragments of *Acropora palmata* on a mild slope, with massive colonies of *Pseudodiploria labyrinthiformis*, *P. clivosa* and *P. strigosa*, *Colpophyllia natans*, *Orbicella annularis* and *O. faveolata*. Scarce soft corals such as *Pseudopterogorgia* sp. and *Gorgonia ventalina*. |
|  | *Orbicella annularis* | Large fused colonies (> 2m height) of *Orbicella annularis* (discreet units cannot be discriminated) with sparse colonies of *Acropora cervicornis*, *Pseudodiploria labyrinthiformis*, *P. clivosa* and *P. strigosa*, *Colpophyllia natans*, *O. faveolata* and *Porites astreoides*. Scarce soft corals such as *Pseudopterogorgia* sp. and *Gorgonia ventalina* and tubular and encrusting sponges. Abundant carbonate macroalgae. |
|  | Octocorals-antipatharian | Large discrete colonies (> 2m height) of *Orbicella annularis* as the main component of the reef frame with sparse colonies of *Acropora cervicornis*, *Pseudodiploria labyrinthiformis*, *P. clivosa* and *P. strigosa*, *Colpophyllia natans*, *O. faveolata* and *Porites astreoides*. Abundant *Pseudopterogorgia* sp., *Gorgonia ventalina* and encrusting and tubular sponges. |
| Pavement | Mounds of *Madracis* sp. | Consolidated mounds (> 30m<sup>2</sup>) of *Madracis* sp. with or without sparse colonies of *Eunicea* sp., *Pseudodiploria clivosa* and *P. labyrinthiformis*. |
|  | Sparse massive colonies | Large (> 1m height) and sparse colonies of *Pseudodiploria labyrinthiformis*, *P. clivosa* and *P. strigosa*, *Colpophyllia natans*, *Siderastrea siderea*, *Orbicella annularis*, *Porites porites* and *Porites astreoides* also carbonate macroalgae such as *Halimeda* sp. and *Turbinaria* sp. |
|  | Octocorals | Dense patches of *Pseudopterogorgia* and *Plexaura flexuosa* with spare and infrequent colonies of *Pseudodiploria strigosa* and *Orbicella annularis*. |

**Table S2.** Descriptive statistic of *Dendrogyra cylindrus* size frequency distribution of colonies within three sand flats (isla Larga, Noronqui and Yonqui), three consolidated (Faro, Cote and Salina) and the MPA.

|  | Faro | Cote | Salina | Isla Larga | Noronqui | Yonqui | MPA |
| --- | --- | --- | --- | --- | --- | --- | --- |
| No. colonies | 10 | 26 | 58 | 25 | 8 | 11 | 198 |
| Min. | 22 | 5 | 15 | 17 | 10 | 17 | 5 |
| Máx. | 260 | 230 | 290 | 86 | 60 | 260 | 280 |
| Average | 136.1 | 79.81 | 75.43 | 49.04 | 46.13 | 60.64 | 76.1 |
| Standard Deviation | 81,49 | 52,88 | 50,24 | 18,51 | 16,01 | 70,77 | 93,84 |
| Variation Components | 59,88 | 66,26 | 66,6 | 37,74 | 34,72 | 98,72 | 123 |
| Median | 126,5 | 65 | 65 | 45 | 51.5 | 31 | 53 |
| Mode | 260 | 50 | 50 | 32 | 60 | 63 | 32 |
| Asymmetry | 0,07 | 0,99 | 1,73 | 0,51 | -1,29 | 1,98 | 6,13 |

